# Cell-free Reconstitution Reveals Synergistic Stabilization of Microtubule Doublets by PACRG and FAP20

**DOI:** 10.1101/2025.03.12.642377

**Authors:** Daria Khuntsariya, Umut Batman, Arya Krishnan, Daniel Rozbesky, Florent Lemaitre, Carsten Janke, Virginie Hamel, Paul Guichard, Marcus Braun, Zdenek Lansky

## Abstract

Ciliary microtubule doublets, consisting of an incomplete B-tubule on the surface of a complete A-microtubule, form the fundamental structural framework of motile and sensory cilia/flagella. Ciliary proteins PACRG and FAP20, which are essential for flagellar stability and function, localize to the junction between the A-and the B-tubule. However, their precise role in microtubule doublet assembly and dynamics remains unknown. Here, using a cell-free assay, TIRF-microscopy, and cryo-electron tomography, we demonstrate that in combination PACRG and FAP20 stabilize B-tubule dynamics and microtubule doublet architecture. We show that together these proteins localize to the B-tubules in distinct, high-density patches, which locally stabilize B-tubule by decreasing its depolymerization velocity and increasing its rescue frequency. Cryo-tomography of *in vitro* reconstructed microtubule doublets in presence of PACRG and FAP20 reveals reduced B-tubule curvature fluctuations, promoting a more rigid and aligned conformation. Altogether, our findings suggest that PACRG and FAP20 function synergistically to reinforce microtubule doublet stability, ensuring ciliary integrity and function.

## Introduction

Cilia are microtubule-based, membrane-bound organelles that project from the surface of eukaryotic cells. Cilia play crucial roles in various physiological processes, including sensing of chemical signals, light, or mechanical stress, as well as enabling cell motility. Dysfunction of cilia can result in sever pathologies collectively termed ciliopathies^1,2^. The core structure of cilia is axoneme. The axoneme architecture has a highly conserved “9+2” arrangement, consisting of nine peripheral microtubule doublets surrounding a central pair of single microtubules. In the case of non-motile, primary cilia, the architecture is composed only of 9 microtubule doublets. In both cases, each microtubule doublet is composed of a complete A-tubule with 13 protofilaments and an incomplete B-tubule with 10 protofilaments, forming a unique structure that is critical for ciliary stability and motility^3–5^.

In cilia, post-translational modifications (PTMs) of tubulin C-terminal tails such as glycylation^6,7^, glutamylation^8^ and different levels of tyrosination and detyrosination^9^ play essential roles in regulating cilia beating^10–13^ or the transport of ciliary components by intraflagellar transport trains^14^. Further, A- and B-tubules undergo differential modifications, with A-tubules being glycylated^15^ and tyrosinated^9^ and B-tubules being highly detyrosinated^9,14^, glycylated and polyglutamylated^11^. Dysregulation of tubulin post-translational modifications is associated with ciliopathies^12^, with polyglutamylation being of particular functional importance^13^. For example, recent studies have shown that excessive glutamylation, implicated in retinitis pigmentosa, alters the molecular architecture of the primary cilium in photoreceptors^16^. Additionally, dysregulation of glutamylation and glycylation is often associated with axoneme defects^17–22^, underscoring their importance in maintaining ciliary integrity.

Electron cryo-microscopy studies of *Chlamydomonas reinhardtii*, as well as mammalian and human microtubule doublets have identified at least 35 microtubule-inner proteins associated with microtubule doublets^23–25^. These proteins are integral to the microtubule structure and ensur proper ciliary function, which includes both motility and signal transduction. However, the functions of the most of these proteins remain unclear.

The interactions between the A-and tubules are mediated by structural junctions, including the outer junction, formed by interactions among protofilaments B1, A10, and A11, and the inner junction, the closure of the B-tubules on protofilaments A1 and A13 (Fig. 1 A). Previously, we demonstrated that the side-to-surface tubulin interaction of the outer junction can be reconstructed *in vitro* by removing the tubulin C-terminal tails of a pre-assembled A-tubule and adding free tubulin^26^. This leads to the formation of a B-tubule on the A-tubule. However, the B-tubule remained open at the inner junction, likely due to the absence of additional proteins required to stabilize the closure. This structural instability results in significant mobility of the “open” B-tubule at the microtubule doublet inner junction^26^. *In vivo*, two non-tubulin proteins, PACRG and FAP20^27,28^, localize to the inner junction, where they are arranged in an alternating pattern along the inner junction, linking protofilament A1 of the A-tubule and protofilament B10 of the B-tubule with an 8-nm periodicity^23^. Based on the structural studies of microtubule doublets, it was shown that PACRG interacts with β-tubulin on the A-tubule, while FAP20 binds to α-tubulin on the A-tubule^23,29^. Both PACRG and FAP20 are highly conserved across ciliated organisms, and their loss is associated with cilia-related phenotypes, such as infertility^30^ or retinal dystrophy^31^, highlighting their critical role in ciliary function. Although FAP20 and PACRG are known to localize at the inner junction and important for cilia motility^27,28^, their precise roles in microtubule doublet assembly, stability, and the mechanism of their insertion into the inner junction are unknown.

**Fig 1.**
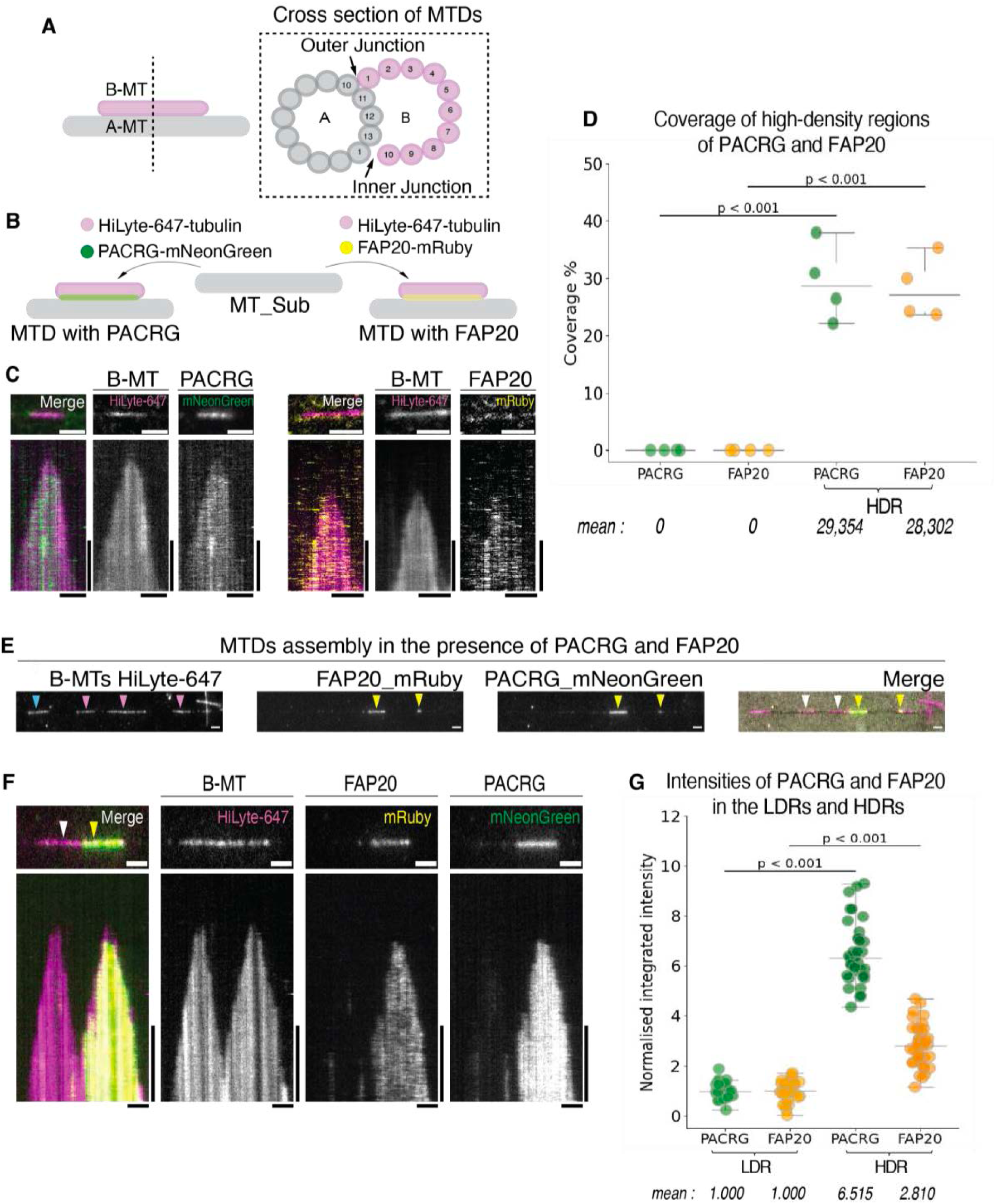
The formation of HDR of PACRG and FAP20 on MTDs i*n vitro*. (**A**) Schematic representation of microtubule doublets (MTDs) cross section, showing outer junction (OJ), inner junction (IJ) and protofilament patterns of A-and B-tubule (A-MT, B-MT). (**B**) Schematic drawing of microtubule doublets (MTDs) assembly in the presence of only one protein, PACRG or FAP20. (**C**) Montage showing B-tubule (B-MT) with binding of PACRG and FAP20, and the corresponding multichannel kymograph. Scale bare, horizontal - 2μm, vertical – 5 min. (**D**) Percentage of high-density regions (HDRs) coverage of PACRG (green) and FAP20 (yellow) along B-tubules length. Groups represent assembled microtubule doublets (MTDs) in the presence of PACRG alone (N=4), FAP20 alone (N=4), and in the presence of PACRG and FAP20 together (N=4), N – number of experiments. p < 0.001, determined by the Mann-Whitney U Test. (**E**) Montage showing assembled microtubule doublets (MTDs) in the presence of 200 nM PACRG-mNeonGreen and 200 nM FAP20-mRuby3 and proteins binding in high-density regions (HDRs) and low-density regions (LDRs). Yellow arrowheads indicate high-density regions (HDRs), white arrowheads indicate low-density regions (LDRs), magenta arrowheads indicate B-tubule (B-MT, depicted in magenta), and blue arrowheads indicate extension of A-tubule (A-MT). Scale bare, 2μm. **(F)** Montage showing B-tubule (B-MTs) with high-density region (HDR) and low-density region (LDR) of PACRG and FAP20 and the corresponding multichannel kymograph. Yellow arrowheads indicate high-density region (HDR), white arrowheads indicate low-density region (LDR). Scale bare, horizontal - 2μm, vertical – 5 min. **(G)** Normalized integrated intensities of low-density regions (LDRs, n=41) and high-density regions (HDRs, n=45) of PACRG and FAP20. p < 0.001, determined by the Mann-Whitney U Test between PACRG (green) of low-vs high-density regions, and respectively FAP20 (orange) of low-vs high-density regions, n – number of analyzed regions.

Here, we show that in combination PACRG and FAP20 can localize to the open B-tubules in a distinct, high-density binding mode. We show that these high-density regions of PACRG and FAP20 stabilize the B-tubule *in vitro* and we reveal the impact of post-translation modification of tubulin C-terminal tail on the binding of PACRG and FAP20. Using cryo-tomography, we further show that the addition of PACRG and FAP20 reduces structural fluctuations of the B-tubule and promotes an apparent closure of the inner junction. Altogether, this work elucidates a potential mechanism by which these two proteins associate with the inner junction of microtubule doublets and highlights the critical roles of PACRG and FAP20 in stabilizing the B-tubule and preventing its depolymerization.

## Results

### PACRG and FAP20 together display two binding modes on microtubule doublets

To assess the role of FAP20 and PACRG in microtubule doublet formation we generated recombinant, fluorescently labelled, full-length, *C. reinhardtii*, PACRG and FAP20 proteins, N-terminally tagged with mNeonGreen and mRuby3, respectively (Methods, Fig. S1 A-C). First, we tested the interaction of PACRG and FAP20 with the A-tubule lattices. To this end we *in vitro* reconstituted GMPCPP-stabilized microtubules, immobilized the microtubules on a coverslip, added either 50 nM PACRG-mNeonGreen, or 50 nM FAP20-mRuby and imaged in TIRF microscopy (Methods). We found that both PACRG and FAP20 bind to microtubules, which strongly suggests that both proteins can directly interact with the A-tubule lattice independently (Fig. S1-D). To study PACRG and FAP20 interactions with B-tubules, we *in vitro* reconstituted open microtubule doublets by adding purified tubulin in presence of GMPCPP to pre-assembled, coverslip-immobilized, subtilisin-treated microtubules (Methods), as previously described^1^. We then supplied the growing microtubule doublets with 200 nM PACRG and 200 nM FAP20, either in combination or both proteins separately. We found that PACRG alone bound to microtubule doublets with higher affinity than to A-microtubule lattices; while FAP20 alone displayed only a slight preference for microtubule doublets (Fig. 1B, C). However, when combined, strikingly, PACRG and FAP20 interacted with microtubule doublets in two distinct ways. Either, they formed prominent, high-density patches, which grew at their edges, suggesting localized cooperative binding (Fig. 1E-G), or, outside of the distinct patches, they displayed modest preference for the microtubule doublets, similar to our observations in presence of only one of the proteins (Fig. S1F). PACRG and FAP20 high-density patches, hence, did not cover all available microtubule doublets. Yet, importantly, we observed the formation of high-density patches of PACRG and FAP20 exclusively on microtubule doublets and only when both proteins were present (Fig. 1D). Further, PACRG and FAP20 high-density patches did not necessarily appear immediately upon microtubule doublet formation, rather they nucleated from diffraction-limited spots on already formed microtubule doublets, with both proteins simultaneously appearing at high density (Fig. 1F). Our data hence indicates that PACRG and FAP20 do not co-assemble with the B-tubules during their polymerization, but rather bind to pre-existing microtubule doublets. To test the hypothesis, we added both proteins together to pre-assembled microtubule doublets and found that PACRG and FAP20 indeed again locally formed high-density patches that did not necessarily cover the whole microtubule doublet length and coexisted with regions of low-density binding of PACRG and FAP20 (Fig. S1 G, H). Higher protein concentrations resulted in increased coverage of the microtubule doublets with high-density patches of PACRG and FAP20. Combined, these results show that when binding together to an open B-tubule, PACRG and FAP20 can form high-density patches, which nucleate and grow at their edges along the B-tubule.

### FAP20 and PACRG are more stable in high-density regions

Given the putative position at the junction between A-and B-tubule lattices, PACRG and FAP20 might be well integrated into the microtubule structure. To test this, we investigated the turnover of PACRG and FAP20 in high-density versus low-density patches. To quantify turnover, we added both proteins together to pre-assembled microtubule doublets and measured the fluorescence recovery after photobleaching (FRAP) in regions of high and low density, respectively (Fig. 2A-B). We found that in low-density binding regions, PACRG and FAP20 signals recovered rapidly and almost completely after bleaching. In contrast, in high-density binding regions, signal recovery was significantly slower and less complete for both proteins, indicating a markedly reduced turnover. Furthermore, our results suggest that under high-density binding conditions, approximately one third of PACRG and FAP20 molecules remain stably bound and do not turn over at all (Fig. 2C-F). Together, these data indicate that PACRG and FAP20 molecules that localize in high-density patches are significantly less mobile compared to their binding outside of these regions, in line with a hypothesis that they might be embedded in the MTD structure.

**Fig 2.**
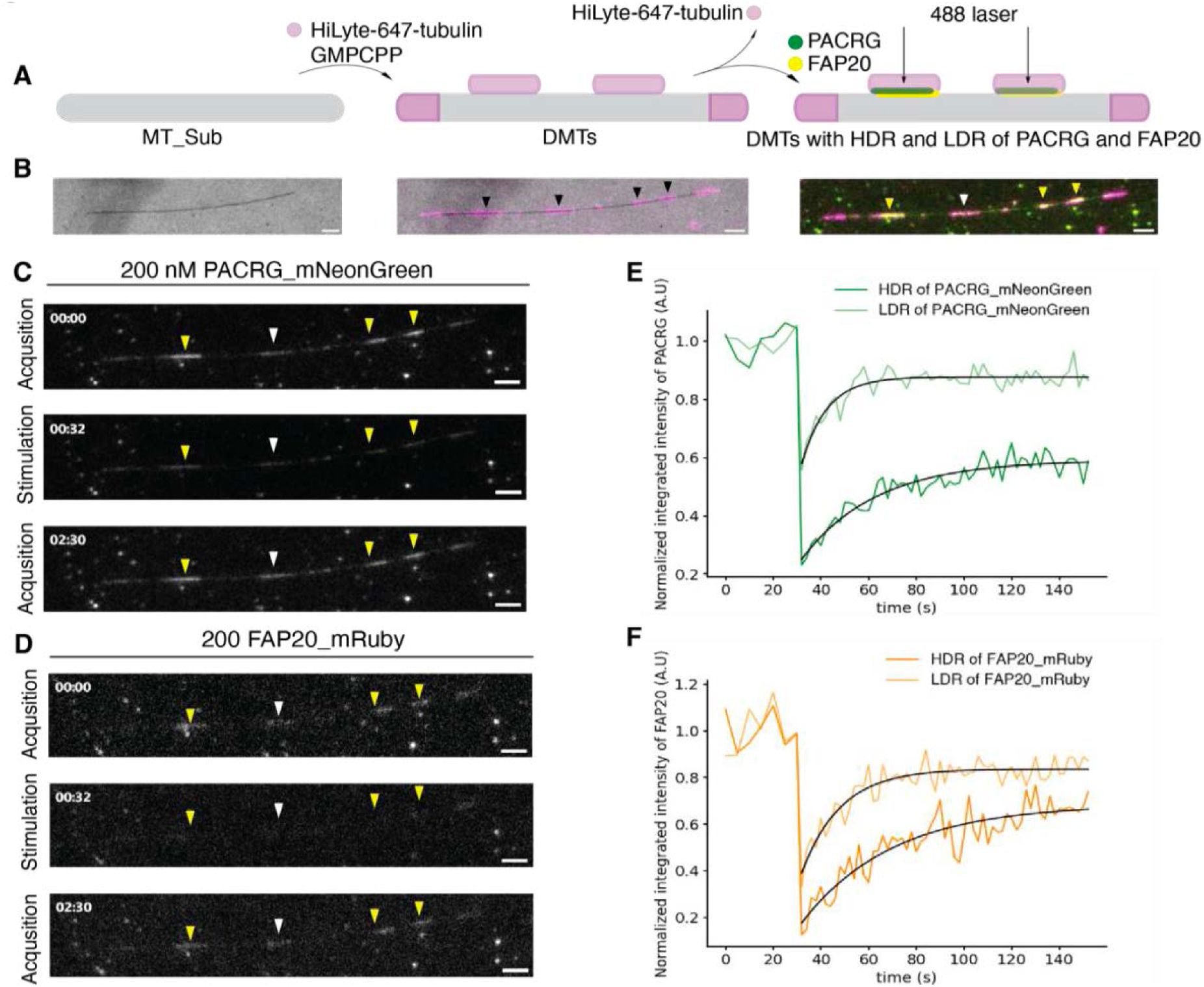
FAP20 and PACRG are more stable in high-density regions. (**A**)Schematic drawing of FRAP assay. (**B**) Montage showing assembled microtubule doublets (MTDs) with high-density regions (HDR) of PACRG and FAP20 before FRAP. B-tubule (B-MT) formed by 647-HiLyte tubulin are depicted in magenta, black arrowheads indicate B-tubule (B-MTs), yellow arrowheads indicate high-density regions (HDRs), white arrowheads indicate low-density region (LDR). Scale bare, 2μm. (**C**-**D**) Montage showing FRAP of PACRG-mNeonGreen (C) and FAP20-mRuby (D) on microtubule doublets (MTDs), yellow arrowheads indicate high-density regions (HDRs), white arrowheads indicate low-density region (LDR). Scale bare, 2μm. (**E**) Normalized integrated intensities of high-density regions (HDRs, dark green) and low-density region (LDRs, light green) during FRAP experiment. The normalized recovery time for PACRG-mNeonGreen in high-density regions (HDR) is 18.618 seconds and in LDR is 4.402 seconds. The immobile fraction was 48% in high-density regions (HDRs) and 17% in low-density regions (LDRs). Number of analyzed FRAP areas for high-density regions (HDRs) n=13 for low-density regions (LDRs) n=17. (**F**) Normalized integrated intensities of high-density regions (HDRs, dark green) and low-density regions (LDRs, light green) during FRAP experiment. The normalized recovery time for FAP20-mRuby in high-density regions (HDRs) is 30.963 seconds and in low-density regions (LDRs) is 16.707 seconds. The immobile fraction was 25% in high-density regions (HDRs) and 11% in low-density regions (LDRs). Number of analyzed FRAP areas for high-density regions (HDRs) n=13 and for low-density regions (LDRs) n=17.

### PACRG and FAP20 stabilize B-tubule

Given the potential positioning of PACRG and FAP20 in the gap between the incomplete B-tubule and the A-tubule lattice^23^, we wondered whether the two proteins might affect B-tubule dynamics. To answer this question, we first examined whether PACRG and FAP20 interact in solution, as suggested by previous studies^27^, and, more importantly, whether they do so independently, or in combination with tubulin. To test this, we mixed His-tagged PACRG with untagged tubulin and/or untagged FAP20 and pulled down the PACRG with NiNTA-beads. When we eluted bound proteins from the beads and analyzed them via SDS-PAGE, we observed that FAP20 and tubulin, both individually and in combination, co-eluted with PACRG. Control experiments showed that FAP20 and tubulin did not interact with beads in the absence of PACRG (Fig 3A). This result indicates that PACRG and FAP20 can directly interact in solution and can form a complex with soluble tubulin.

**Fig 3.**
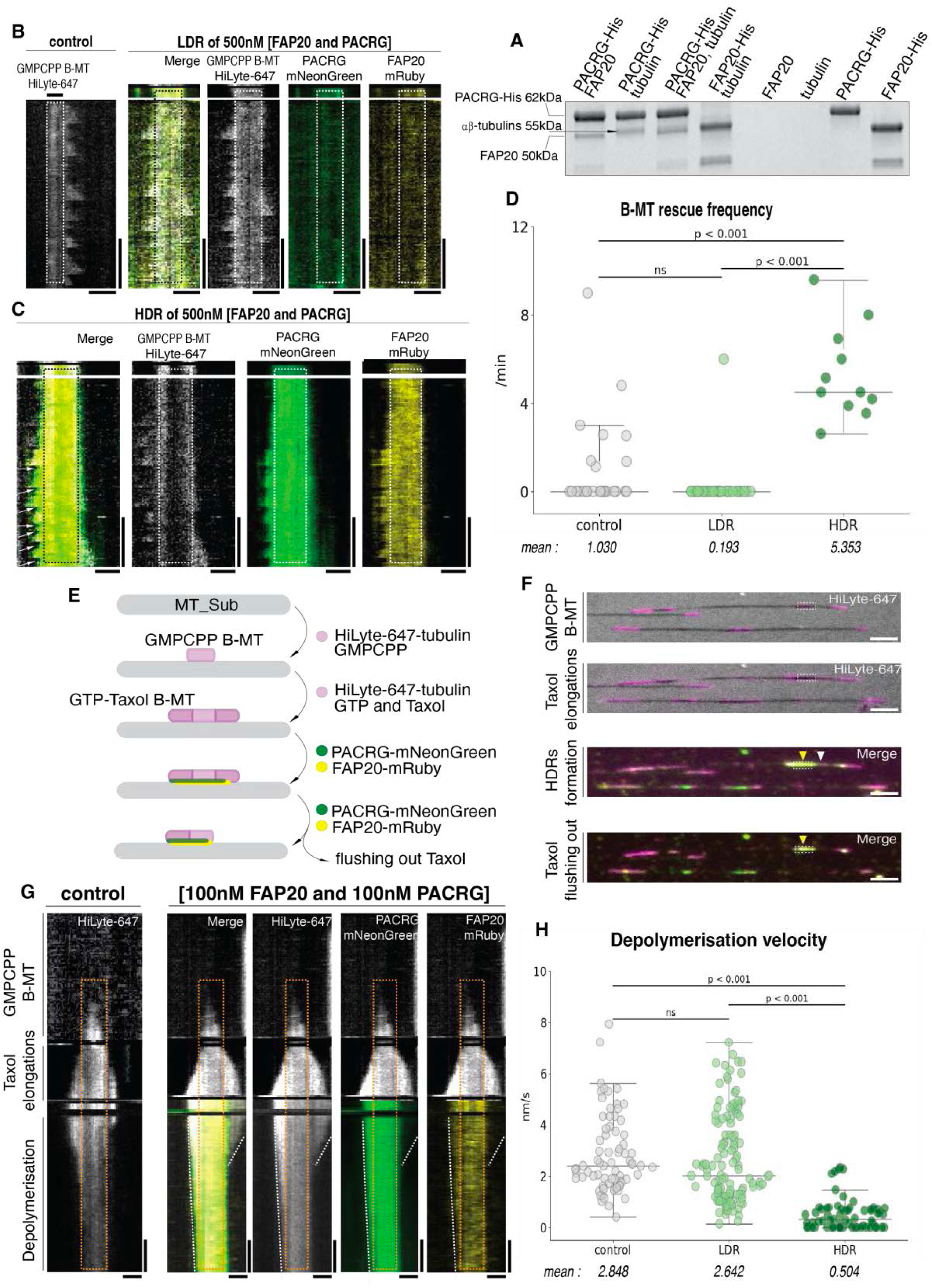
PACRG and FAP20 stabilize B-tubule. (**A**) SDS-PAGE gel of pull-down assay showing the elution fraction of the proteins bonded to the beads. Mw of PACRG-His is 62 kDa, FAP20 is 49.7 kDa, FAP20-His is 50.5 kDa and αβ-tubulins 55 kDa. **(B-C)** Montage showing dynamic instability of B-tubules (B-MTs) without addition of proteins and with accumulation of both proteins (500nM PACRG-mNeonGreen and 500 nM FAP20-mRuby3) in low-density region (LDR) on B-tubules (B-MTs) (B) and with high-density region (HDR) on B-tubule (B-MT) (C). Dashed rectangles indicate GMPCPP seeds of B-tubule (B-MT). White arrowheads indicate rescues. Horizontal scale bar 1 μm, Vertical scale bar 5 min. (**D**) B-tubule (B-MT) rescue frequency (min^-1^) of B-tubule (B-MTs) without addition of proteins (control), B-tubule (B-MTs) with low-density binding of proteins (LDR), and B-MTs with high-density binding of proteins (HDR). Means and p-value were determined by two-tailed student test. Analyzed number of B-tubule (B-MTs): control n= 25, LDR n=30, HDR n= 11). (**E**) Schematic drawing of depolymerization assay. (**F**) Montage showing snapshots of depolymerization assay: formation of Taxol-GTP B-tubule (B-MTs) elongations from GMPCPP-B-seeds and further depolymerization in the presence of 100 nM PACRG-mNeonGreen and 100 nM FAP20-mRuby3. White dashed rectangles indicate GMPCPP-B-tubule. Yellow arrowheads indicate high-density region (HDR), white arrowheads indicate low-density region (LDR). Scale bare, 2μm. (**G**) Montage showing corresponding multichannel kymographs of depolymerization assay with and without 100 nM PACRG and 100 nM FAP20 together. Orange dashed rectangles indicate preassembled GMPCCP-B-tubule seeds. White dashed line indicates depolymerization of B-tubule (B-MTs) with low-density bindings of proteins. Horizontal scale bar 1 μm, Vertical scale bar 5 min. **(H)** Depolymerization velocity (nm/s) of GDP-taxol-B-tubules elongations. Means and p-value were determined by two-tailed student test. Analyzed number of B-tubules: control n= 36, LDR n=62, HDR n= 32.

To explore how FAP20 and PACRG affect the dynamics of B-tubules, we added free tubulin - in presence of GTP - to preassembled microtubules doublets, both in presence and absence of PACRG and FAP20 (Methods). For comparison, we analyzed in parallel the dynamics of A-tubules and B-tubules, which we acquired in the same imaging experiments. We found that similar to A-tubules, B-tubules are more dynamic at their plus ends then the minus ends (Fig. S3 B-D), displaying a higher frequency of catastrophes (Fig. S3F). Strikingly, while the addition of 500 nM FAP20 and 500 nM PACRG together had no significant effect on A-tubules dynamics (Fig. S3J-N), high-density patches of both proteins on B-tubules strongly increased the rescue frequency of B-tubules (Fig. 3B-D). Additionally, growth velocity and catastrophe frequency of B-tubules were modestly reduced by the presence of high-density patches of PACRG and FAP20 (Fig. S3H,I). In contrast, in regions of low-density binding of PACRG and FAP20 the dynamic parameters of B-tubules were not affected (Fig. 3B,D; S3H,I). Together, our data demonstrate that high-density PACRG and FAP20 patches with low-turnover specifically stabilize B-tubules.

During our experiments, we noticed that B-tubule shrinking velocities tended to be reduced in regions of high-density FAP20 and PACRG binding. However, the limited temporal resolution of our imaging setup did not enable reliable quantification of the shrinking velocities. To overcome this limitation, we employed the microtubule-stabilizing drug taxol, which reversibly stabilizes microtubules and enables synchronized depolymerization upon its removal, allowing for better measurement of depolymerization velocity. For this assay, we first generated short B-tubule seeds using GMPCPP, then elongated them with free tubulin, GTP, and taxol. This resulted in B-tubule extensions composed of GDP-tubulin, which are normally prone to depolymerization, but were transiently stabilized by taxol. Subsequently, we added PACRG and FAP20, which formed high-density patches on the B-tubules on both the irreversibly stabilized GMPCPP sections and the reversibly stabilized GDP-taxol-sections. Next, by flushing the assay chamber with fresh buffer to remove taxol while keeping PACRG and FAP20 in solution, allowing us to monitor the depolymerization of B-tubule GDP sections (Fig. 3E-G). Measuring the depolymerization velocity, we found that in regions of low-density PACRG and FAP20 binding, depolymerization velocities were indistinguishable from those of control B-tubules lacking additional proteins. By contrast, at high-density PACRG and FAP20 regions, B-tubules exhibited a markedly slower depolymerization rate (Fig. 3H). Combined, these experiments demonstrate that PACRG and FAP20 increase the stability of B-tubules.

### Tubulin posttranslational modifications regulate PACRG and FAP20 interaction with microtubules

Tubulin post-translational modifications are important for the function of cilia. We thus aimed to determine whether PACRG and FAP20 binding to microtubules is influenced by post-translational modifications. To do this we employed microtubules assembled from tubulin derived from different sources. This included HeLa cells, which provide tyrosinated tubulin lacking other modifications^32^ (Methods) and wild type mouse brain tissue, providing highly modified tubulin^33^ (Methods). Additionally, we used tubulin from knockout mouse models deficient in specific tubulin-modifying enzymes, resulting in tubulin specifically lacking polyglutamylation on α-tubulin (*Ttll1^−/−^*), β-tubulin (*Ttll7^−/−^*), or both (*Ttll1*^−/−^*Ttll7*^−/−^), as well as tubulin lacking acetylation on α-tubulin (*Atat1^−/−^*, Methods)^34,35^ Using TIRF microscopy, we first compared the binding of PACRG and FAP20 to highly modified (mouse brain) vs. unmodified (HeLa) microtubules to test whether the proteins are affected at all by post-translational modification.

To ensure identical experimental conditions, we used unlabelled tubulin and imaged two different types of microtubules in the same assay chamber. To distinguish the microtubule subtypes, we first immobilized one type on a coverslip, imaged their positions, and then added the second type of microtubules. We subsequently added the fluorescently labelled proteins and quantified their normalized integrated intensities on the two different types of microtubules. Our analyses revealed a preference of PACRG-mNeonGreen for wild-type mouse brain microtubules over HeLa microtubules, while FAP20 did not show any preferential binding (Fig. S4A-E). This suggested that PACRG is sensitive to one of the post-translational modifications present on brain tubulin. To determine which modification affects PARRG binding, we further compared its binding affinities to wild-type mouse brain microtubules, which are highly polyglutamylated and acetylated, compared to microtubules from our knockout mouse models that lack subsets of these PTMs. We found that PACRG displays the highest affinity for modified (wild type) microtubules, while absence of polyglutamylation on either α-(*Ttll1^−/−^*) or β-tubulin (*Ttll7^−/−^*) reduces the affinity. Moreover, we found that this effect was roughly additive, with the double knockout (*Ttll1^−/−^Ttll7^−/−^*) displaying the strongest reduction. By contrast, acetylation had no impact on PACRG binding, as binding to wild-type and *Atat1^-/-^* microtubules was similar (Fig. 4A-C). Finally, we investigated the role of detyrosination by comparing PACRG and FAP20 binding to microtubules assembled from HeLa tubulin, which is natively predominantly tyrosinated, or HeLa microtubules enzymatically detyrosinated with Carboxypeptidase A^36^ (Methods). We found that both PACRG and FAP20 preferentially bind to tyrosinated microtubules (Fig. 4D-H). Combined, these experiments demonstrate that PACRG binds preferentially to polyglutamylated microtubules, while both FAP20 and PACRG preferentially bind to tyrosinated microtubules.

**Fig 4.**
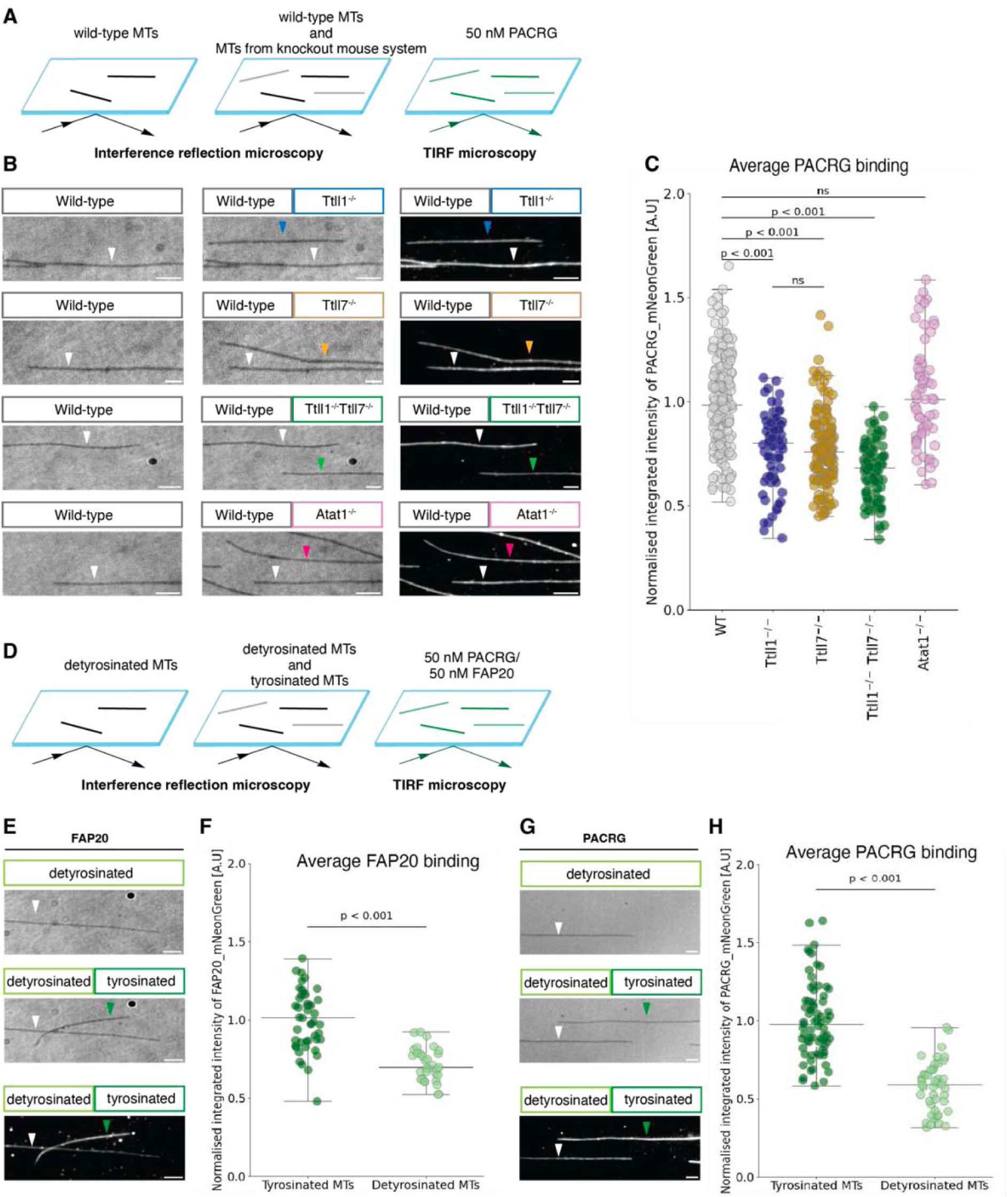
PTMs of a C-terminal tail of tubulin affects binding of PACRG and FAP20 to microtubules. (**A**) Schematic drawing of the assay setup. (**B**) Montage showing IRM and TIRF microscopy images of the microtubules polymerized from wild-type and knockout-mouse brain tubulin and 50 nM PACRG-mNeonGreen intensities for different post-translational modification (PTM) variant. White arrowheads indicate wild-type, blue - *Ttll1^−/−^*, orange - *Ttll7^−/−^*, green - *Ttll1^−/−^Ttll7^−/−^*, pink – *Atat1^−/−^*. Scale bar, 2μm. (**C**) Normalized integrated average intensities of PACRG-mNeonGreen on Taxol-MTs, in three independent sets of the experiment, N=3. P-values, determined by the Mann-Whitney U Test between wild type (WT) and each post-translational modification (PTM) variant and between *Ttll1^−/−^* and *Ttll7^−/−^*. (**D**) Schematic drawing of the assay setup. (**E, G**) Montage showing IRM and TIRF microscopy images of the detyrosinated and tyrosinated microtubules and intensities of FAP20-mNeonGreen and PACRG-mNeonGreen respectively. White arrowheads indicate detyrosinated microtubules, green arrowheads - tyrosinated microtubules. Scale bar, 2μm. (**F, H**) Normalized integrated average intensities of PACRG-mNeonGreen and FAP20-mNeonGreen respectively on Taxol-microtubules, in three independent sets of the experiment, N=3. P-values, determined by the Mann-Whitney U Test.

### FAP20-PACRG Stabilizes B-Microtubule Curvature Within Microtubule Doublets

To investigate the role of PACRG and FAP20 in microtubule doublet closure and shape we used cryo-electron tomography (cryo-ET) to visualize the architecture of *in vitro* assembled microtubule doublets, either in the absence or presence of PACRG and FAP20 (Fig. 5). Our cryo-ET reconstructions offer a detailed and comprehensive view of the structural organization of microtubule doublets *in vitro*. To preserve the intrinsic structural heterogeneity, which is often lost through averaging, we deliberately chose not to apply this technique to our tomograms as previously^26^. This allowed us to capture the full range of natural variations within the microtubule doublet architecture assembled *in vitro*, providing insights that might otherwise be overlooked in traditional averaging-based analyses.

**Fig. 5.**
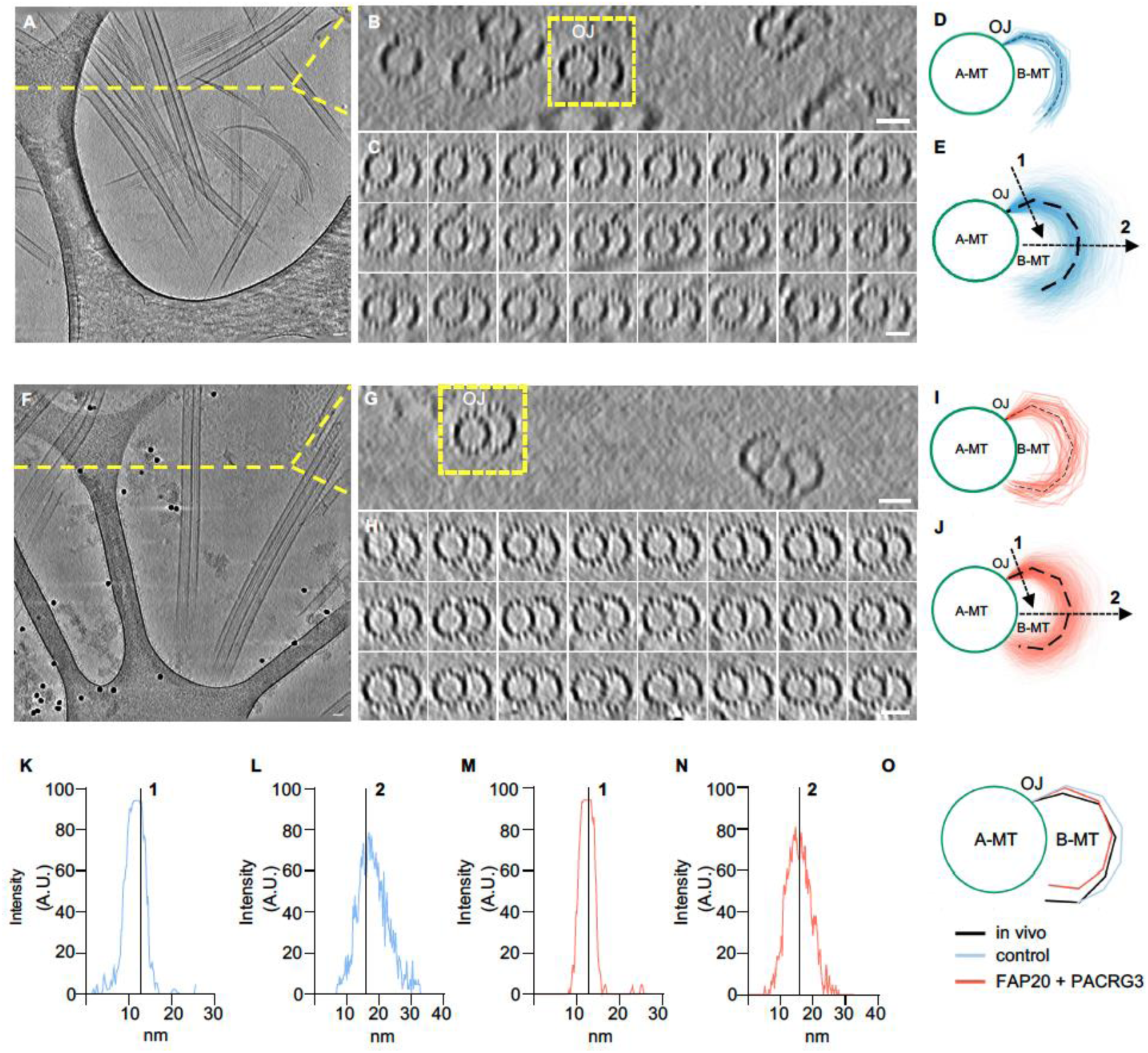
FAP20-PACRG Stabilizes B-Microtubule Curvature Within Microtubule Doublets. **(A**) Representative image of a cryo-ET section of microtubule doublets (MTDs) alone. Scale bar, 25 nm. (**B**) *zx* view of a cryo-ET section. Tomogram in (A) is resliced from the top and projected with average intensity using grouped z-projection with a group size of 10 slices corresponding to 8 nm thickness. Scale bar, 25 nm. (**C**) Montage of microtubule doublet (MTD) shown in (B). Cross section of grouped z-projection along the microtubule doublet (MTD) length (376 nm) with 8 nm spacing. Scale bar, 25 nm. (**D**) Coordinates of the B-tubule in (B) starting at the OJ for the microtubule doublet (MTD). Black dashed line indicates the average (**E**) Coordinates of all B-tubules starting at the OJ for *in vitro* MTDs alone (n=46 microtubules from 23 tomograms). Black dashed line indicates the average. (**F**) Representative image of cryo-ET section of microtubule doublets (MTDs) in the presence of FAP20-PACRG. Scale bar, 25 nm. (**G**) *zx* view of a cryo-ET section. Tomogram in (F) is resliced from the top and projected with average intensity using grouped z-projection with a group size of 10 stacks corresponding to 8 nm thickness. Scale bar, 25 nm. (**H**) Montage of microtubule doublet (MTD) shown in (G). Cross section of grouped z-projection along the microtubule doublet (MTD) length (560 nm) with 16 nm spacing. Scale bar, 25 nm. (**I**) Coordinates of the B-tubule in (G) starting at the outer junction (OJ) for the microtubule doublet (MTD). Black dashed line indicates the average. (**J**) Coordinates of all B-tubules starting at the outer junction (OJ) for in vitro microtubule doublets (MTDs) in the presence of FAP20-PACRG (n=40 microtubules from 22 tomograms). Black dashed line indicates the average. (**K** and **L**) Plot profiles at the positions indicated by the arrows in (E) showing the curvature flexibility of B-tubule of microtubule doublets (MTDs) in the absence of FAP20-PACRG. The black line indicates the position of the B-tubule *in vivo*. A.U., arbitrary units. (**M** and **N**) Plot profiles at the positions indicated by the arrows in (J) showing the curvature flexibility of B-tubule of microtubule doublets (MTDs) in the presence of FAP20-PACRG. The black line indicates the position of the B-tubule *in vivo*. A.U., arbitrary units. (**O**) Comparison of average B-tubule curvatures of *in vivo*, microtubule doublet (MTDs) alone and microtubule doublet (MTDs) in the presence of FAP20-PACRG.

In control conditions, where microtubule doublets are assembled without additional proteins, we found that B-tubules do not form the closed inner junction with the A-tubules and display significant curvature fluctuations along their length (Fig. 5A-E, K-L). This intrinsic flexibility likely reflects the absence of stabilizing interactions within the doublet structure, leading to deviations from the more constrained B-tubule conformation observed in native axonemal doublets. Upon the addition of FAP20-PACRG, we observed a striking stabilization of B-tubule curvature, with reduced deviations from the *in vivo* reference position and an apparent closure of the inner junction (Fig. 5F-J, M-N). The plot profiles of individual microtubules show a marked decrease in curvature variability, indicating that FAP20-PACRG constrains B-tubule movement relative to the A-tubule (Fig. 5O). This effect is consistent across multiple tomograms, suggesting that FAP20-PACRG directly reinforces B-tubule architecture by promoting a more rigid and aligned conformation.

## Discussion

Here, we show that the ciliary proteins PACRG and FAP20 play a synergistic role in the formation and stabilization of microtubule doublets. To demonstrate this, we used cell-free *in vitro* reconstitution of microtubule doublets which allow the formation of open and flexible B-tubule. Our findings indicate that PACRG together with FAP20 can co-localize to microtubule doublets in two distinct population: high-density patches and low-density binding regions. High density patches assemble exclusively in the presence of both proteins, PACRG and FAP20. Using dynamic assay of B-tubule we show that high density patches of PACRG and FAP20 stabilize microtubules by significantly increasing rescue frequency and reducing depolymerization speed. *In vivo* microtubule doublets are characterized by reduced dynamic instability, which is essential for maintaining the structural integrity and ciliary function^37,38^. Our findings demonstrate that interplay of both, PACRG and FAP20 proteins might be essential for the microtubule doublet stability regulation. The interplay between these two proteins is further corroborated by in vivo studies in *C. reinhardtii*, where the FAP20 knockdown resulted in reduced endogenous level of PACRG^28^ and exogenous addition of both PACRG and FAP20 together to the *C. reinhardtii* FAP20:PACRG double knockout mutant was more efficient compared to the addition of only one of the proteins^27^. Finally, cryo-electron tomography of *in vitro* assembled microtubule doublets in the presence of PACRG and FAP20 further support the notion that FAP20-PACRG functions as a structural stabilizer within MTDs and suggests that these two proteins may be sufficient to close the inner junction between A-and B-tubule.

We show that PACRG and FAP20 together with soluble tubulin can form a complex independent of the microtubule lattice. This finding suggests the possibility that PACRG and FAP20 may form a complex with tubulin before assembling onto microtubule doublets. Our data however do not favour a model where PACRG and FAP20 co-assemble with the B-tubule during its polymerization. They rather favour a model in which inner junction formation starts from an assembly of an open and flexible B-tubule of 10 protofilament, and subsequent bridging by PACRG and FAP20 and closure of the inner junction. We noted that in the current study we observed that B-tubules in the presence of GTP and tubulin exhibit dynamics preferentially on the plus end (Fig S3 B2-C3). However in the presence of stabilization agents such as GMPCPP (Fig. 1B,E) and taxol (Fig. 3F), B-tubules grow isotropically, as was previously described^26^.

The tubulin C-terminal tails play a crucial role in the *in vivo* microtubule doublet assembly and display an uneven surface distribution across the doublet structure, A-tubule is known to be rather little posttranslationally modified, which includes high levels of tyrosination, given that most α-tubulin isotypes carry a gene-encoded tyrosine. By contrast, the B-tubule exhibit high level of a variety of PTMs, including polyglycylation and polyglutamylation^9,39^. A previous studies showed that C-terminal tails of β-tubulin on protofilament 1 of A-tubule bind the interface between PACRG and FAP20^23,29^. We demonstrate here that both PACRG and FAP20 can bind preferentially tyrosinated microtubules, moreover PACRG bind preferentially to polyglutamylated microtubules, suggesting that these PTMs might be factors that modulate PACRG and FAP20 recruitment to the microtubules and thus the microtubule doublet formation.

Our results suggest that FAP20-PACRG together function as a structural stabilizer within microtubule doublets, potentially reinforcing microtubule doublet integrity in cilia and flagella. By limiting B-tubule flexibility, FAP20-PACRG may contribute to the mechanical resilience required for motile and sensory functions. Further investigations should dissect how this stabilization influences microtubule doublet assembly and its interplay with other microtubule-associated proteins in vivo.

## Acknowledgements

This work was supported the Swiss State Secretariat for Education, Research and Innovation (SERI) under contract number MB22.00075 attributed to P.G., the ISREC TANDEM attributed to V.H, and the Swiss National Foundation 310030_205087 attributed to P.G. and V.H, by Charles University Grant Agency (GAUK 165523) to D.K, by Czech Science Foundation (grant 23-07703S to M.B. and Junior Star Grant 21-27204M to D.R.), the European Research Council (grant ERC-2022-SYG 101071583) to CJ and ZL, the Agence Nationale de la Recherche grant ANR-20-CE13-0011 to CJ, the Fondation pour la Recherche Médicale grant FDT202304016528 to AK, and institutional support from the CAS (RVO: 86652036). We thank Katerina Konecna, Tereza Smidova (IBT, Prague), and Laura Lebrun (Institut Curie) for technical support. We acknowledge the Imaging Methods Core Facility at BIOCEV, institution supported by the MEYS CR (LM2023050 Czech-BioImaging), the CF Protein Production of CIISB, Instruct-CZ Centre, supported by MEYS CR (LM2023042).

## Autor contribution

Conceptualization, M.B, and Z.L; methodology, D.K., U.B., A.K., C.J., V.H, P.G, M.B., Z.L.; investigation, D.K., U.B, A.K; formal analysis, D.K., U.B., F.L.; data curation D.K., U.B.; recourses; A.K., C.J., D.K., U.B., D.R. writing, D.K.,V.H., P.G., M.B., Z.L.; visualization, D.K., U.B.; supervision, M.B., Z.L., V.H., P.G.; finding acquisition, D.K., C.J., V.H., P.G., M.B., Z.L.

## Methods

### Purification of recombinant FAP20-mRuby and PACRG-mNeonGreen

Full-length crPACRG and crFAP20 were isolated and amplified from PACRG cDNA clone in pET30a^2^ FAP20 cDNA clone in pET30a^2^. The nucleotide sequences were inserted to destination vectors congaing C3-cleavage site and fluorescent tags mNeonGreen and mRuby at the N-terminus of PACRG and FAP20 respectively. Both constructs were expressed in insect cells using DefBac viral backbone using FlexiBac Protocol^8^. 72 hours post transfection, the cells were harvested by at 4°C for 10 min at 300 x g. The cell pellets was resuspended in cold lysis buffer (50 mM Na-Phosphate buffer, 300 mM KCl, 1 mM MgCl_2_, 0.1% Tween20, 5% glycerol, 10mM BME, 0.1 mM ATP, 10 mM imidazole, pH 7.5) supplemented with benzonase (1 U ml^−1^; Merck, Darmstadt, Germany) and protease inhibitor cocktail (cOmplete, EDTA free, Roche, Basel, Switzerland) and cells were disrupted by centrifugation at 4°C for 1 hour at 70000 x g. The supernatants were loaded onto a HisPur Ni-NTA Superflow Agarose resin (Thermo Scientific™) equilibrated in lysis buffer for affinity chromatography. After 2 hours incubating the resins with proteins the column was washed by washing buffer (50 mM Na-Phosphate buffer, 300 mM KCl, 1 mM MgCl_2_, 0.1% Tween20, 5% glycerol, 10mM BME, 0.1 mM ATP, 20 mM imidazole, pH 7.5) two times. The proteins were eluted by cleaving off the N-terminal tag with C3 protease in the wash buffer overnight for PACRG and 3 hours for FAP20 at 4°C. Proteins were collected and concentrated using an Amicon ultracentrifuge filter with a molecular weight cutoff of 50 kDa (Merck, Darmstadt, Germany) and loaded onto Superdex 200 10/300 GL column for further separation by size exclusion chromatography with 100 mM Tris-HCl pH 8.0, 150 mM NaCl, 1 mM EDTA, 2 mM MgCl, 0.1% Tween20, 5% glycerol, 1 mM DTT, 1 mM ATP as a mobile phase. The success of purifications was verified by SDS-PAGE (Fig. S1). The purified proteins were concentrated using a Amicon ultracentrifuge filter and freeze using liquid nitrogen.

#### 3.3.1 Pull-down assay

Pull-down assay was performed using the The Thermo Scientific^TM^ Pierce^TM^ Pull-Down PolyHis Protein:Protein Interaction Kit (Thermo Scientific, 21277) following the provided protocol. The 50μl of HisPur Ni-NTA Superflow Agarose (Thermo Scientific, 25215) was used as the agarose bead matrix. The wash buffer contained 10mM imidazole (BRB80, 0.5 mM GTP, 10 mM imidazole, pH 7.4), the elution buffer contained 300 mM imidazole (BRB80, 0.5 mM GTP, 300 mM imidazole, pH 7.4). Bait proteins tagged with 6xHis (three samples obtained PACRG-mNeonGreen-6xHis and one had FAP20-mRuby3-6xHis as Bait protein), were immobilized on the beads and washed with wash buffer to remove unbound proteins. Prey protein candidates (FAP20-mRuby3; unlabeled pig brain tubulin; FAP20-mRuby3 and unlabeled pig brain tubulin; unlabeled pig brain tubulin) were introduced into the column, incubated, and washed to eliminate non-specific interactions. Finally, bound proteins were eluted using elution buffer, and interactions were analyzed via SDS-PAGE to identify protein:protein interactions. To ensure specificity, each protein was also immobilized separately on beads as a negative control (FAP20-mRuby3; unlabeled pig brain tubulin; PACRG-mNeonGreen-6xHis; FAP20-mRuby3-6xHis), allowing the identification and elimination of false positives caused by non-non-specific binding to the resin. The final amount of each used protein was 150 μg.

### Microtubules

Porcine brain tubulin was isolated using the high-molarity PIPES procedure^40,41^.

Biotin-labeled tubulin as well as HiLyte647-labeled tubulin were purchased from Cytoskeleton Inc. (T333P and TL670M, respectively).

*Mouse wild-type brain* tubulin as well as brain tubulin from knockout mouse system (*Ttll1^-/-^, Ttll7^-/-^, Ttll1^-/-^Ttll7^-/-^, Atat1^-/-^*)was purified from mouse brains via cycles of temperature-dependent microtubule polymerization and depolymerization as described in detail previously^34,35^. Briefly, brains were extracted from the skulls of mice immediately after sacrificing the animals. For 1 g of brain tissue, 2 ml of ice-cold lysis buffer consisting of BRB80 (80 mM K-PIPES pH 6.8, 1 mM K-EGTA, 1 mM MgCl2, 1 mM β-mercaptoethanol, 1 mM PMSF, and 1× protease inhibitor cocktail composed of 20 μg/ml leupeptin, 20 μg/ml aprotinin and 20 μg/ml 4-(2-aminoethyl)-benzenesulfonyl fluoride; Sigma-Aldrich) were added, and brains were homogenized using an Ultra-Turrax® blender. Lysates were cleared by centrifugation at 50,000 rpm in TLA-55 fixed-angle rotor (∼112,000 × g; Beckman Coulter) for 30 min at 4°C, from now on referred to as “cold centrifugation”. This first supernatant (later referred to as “SN1”) was adjusted to 1 mM GTP and 1/3 volume pre-warmed 100% glycerol, which after careful mixing was incubated for 20 min at 30°C for microtubule polymerization. Polymerized microtubules were pelleted for 30 min at 112,000 × g, 30°C, referred to as “warm centrifugation”. The pellet was resuspended in BRB80 (1/10 of the initial SN1-volume), and microtubules were disassembled by incubation on ice for 20 min, and occasionally pipetting up-and-down to accelerate their depolymerization.

Solubilized tubulin was cleared via cold centrifugation and the supernatant SN3 was adjusted to final concentrations of 1 mM GTP and 0.5 M PIPES and complemented with 1/3 volume pre-heated 100% glycerol. After careful mixing and incubation for 20 min at 30°C the resulting microtubules were pelleted via warm centrifugation. Microtubules polymerized at high molarity do not contain associated proteins. The pellet was resuspended in 1 × BRB80 (1/40 of the initial SN1-volume), depolymerized on ice for 20 min, and cleared via cold centrifugation. The resulting SN5 was adjusted to 1 mM GTP, supplemented with 1/3 volume of pre-heated 100% glycerol, and incubated for 20 min at 30°C. Microtubules were pelleted by warm centrifugation and resuspended in ice-cold BRB80 (1/40 of the initial SN1-volume). After 15 min on ice, the soluble tubulin was cleared by a final cold centrifugation. The tubulin yield was estimated with a NanoDrop ND-1000 spectrophotometer (Thermo Scientific; absorbance at 280 nm; MW = 110 kDa; ε = 115,000 M−1/cm−1). Samples were adjusted to 4 mg/ml, aliquoted, snap-frozen in liquid N2 and stored at −80°C. Samples for SDS-PAGE were collected and analyzed for each purification cycle step.

### HeLa WT and detyrosinated tubulin purification

Tyrosinated and detyrosinated tubulin was isolated from HeLa S3 cells maintained in suspension culture. These cells were cultivated in spinner bottles containing 1 liter of DMEM supplemented with 10% fetal bovine serum (100 mL), 200 mM L-glutamine (10 mL from a 2 M stock), and 1× penicillin-streptomycin (10 mL from a 100× stock). The cultures were kept on a stirrer table within a cell culture incubator for one week^35^.

After the incubation period, the cell suspension was transferred into 1-liter centrifuge bottles and spun at 250 × g for 15 minutes at room temperature. Then resuspended pellet in a minimal volume of PBS and transferred to a 50 mL Falcon tube, followed by another centrifugation at 250 × g for 8 minutes at 4°C. The supernatant was discarded and added equal volume of ice-cold lysis buffer. Lysed cells using a French press at medium pressure. The lysate underwent centrifugation at 112,000 × g for 30 minutes at 4°C to yield the first supernatant (SN1). To SN1, 0.2 M GTP (1/200 of SN1 volume) and prewarmed 100% glycerol (½ of SN1 volume) were added, initiating polymerization as previously described. The polymerized tubulin (P2) was depolymerized using 1/60 of the SN1 volume of BRB80 buffer. Following another centrifugation, the resulting supernatant (SN3) was collected in an Eppendorf tube for Carboxypeptidase A (CPA) treatment. SN3 was divided into two equal parts (+CPA and - CPA), with 1/300 volume of CPA (Sigma C9268) added to one sample. Both were incubated at 30°C for 5 minutes.

A second round of polymerization was carried out using the same conditions, followed by depolymerization, where 1/100 volume of SN1 was added. This process mirrored the protocol used for tubulin extraction from mouse brains. The third polymerization step was identical for both samples. Finally, the depolymerized tubulin was treated with 1/300 volume of SN1 before dilution and storage, following the previously outlined procedures

***GMPCPP-stabilized biotin-labelled microtubules*** (GMPCPP polymerized) were polymerized from 4 mg/ml porcine brain tubulin (wild type) for 1 hour at 37°C in BRB80 supplemented with 2.7 mM GMPCPP (Jena Bioscience, NU-405). The polymerized microtubules were centrifuged for 30 min at 18000 x g in a Microfuge 18 Centrifuge (Beckman Coulter). After centrifugation the pellet was resuspended in BRB80 at RT.

***Taxol-stabilized microtubules*** were polymerized by adding 1.25 μl of polymerization mix (5mM GTP, 20 mM MgCl_2_, 25% DMSO in BRB80) to 5 μl of 4 mg/ml tubulin (HeLa tubulin, wild-type mouse brain tubulin or mouse brain tubulin with knockout of PTM) for 30 min at 37°C. Subsequently, 200 μl of BRB80 supplemented with 10 μM Paclitaxel was added to microtubules and incubated for 1 min at 37°C and followed by centrifugation (30 min at 18000 x g). the pellet was resuspended in 100 μl of BRB80 supplemented with 10 μM Paclitaxel.

***Subtilisin-treated microtubules*** were polymerized from 4 mg/ml biotin-labelled tubulin (un-labeled tubulin 82%, biotin-tubulin 12%) for 1 h at 37°C in BRB80 supplemented with 2.7 mM GMPCPP (Jena Bioscience, NU-405). The polymerized microtubules were centrifuged for 30 min at 18000 x g in a Microfuge 18 Centrifuge (Beckman Coulter). After centrifugation the pellet was resuspended in BRB80 and microtubules were subsequently treated with subtilisin (CAS Number 9014-01-1, P8038) (1 mg/ml) in a subtilisin:tubulin final ratios of 1:20 (w/w) for 30 min at 30°C. After treatment, subtilisin activity was inhibited by the addition of PMSF (phenylmethylsulphonyl fluorid, 78830) to final concentration 10mM. After incubation at RT for 10 min, subtilisin-treated microtubules (MT_Sub) were next centrifuged at 18,000 x g for 15 min in a Microfuge 18 Centrifuge (Beckman Coulter) to remove both unpolymerized tubulin and subtilisin. The pellet was resuspended in 100 μl BRB80, RT. Subtilisin-treated microtubules were used for microtubule doublet assembly assay^1^.

### TIRF microscopy

Total internal reflection fluorescence (TIRF) microscopy experiments were performed on an inverted microscope (Nikon-Ti E, Nikon-Ti2, Nikon-Ti2-ring/FRAP) equipped with 60x or 100x NA 1.49 oil immersion objectives (Apo TIRF or SR Apo TIRF, Nikon) and either with Hamamatsu Orca Flash 4.0 LT (Hamamatsu Photonics) or PRIME BSI (Teledyne Photometrics) sCMOS cameras. An additional 1.5x magnifying tube lens was used. Microtubules were visualized by using epifluorescence lamp and PACRG-mNeonGreen with FAP20-mRuby or FAP20-nMeonGreen and HiLyte-647 tubulin were visualized sequentially by switching between microscope filter cubes for GFP, mCherry and Cy5 channels. The imaging setup was controlled by NIS Elements software (Nikon).

Flow chambers for TIRF imaging assays were prepared as described previously^42^. Channels were treated with 6 ug/ml anti-biotin solution (Sigma, B3640, 1 mg/ml in PBS) or alternatively with 20 μg/ml anti-β-tubulin (Sigma, T7816-2ML) solution after 5 minutes incubation, followed by one-hour incubation with 1% Pluronic F127 (Sigma, P2443).

### In vitro microtubule doublet assembly assay

Microtubule doublets (MTD) were assembled by adding 8 μM of HiLyte647-tubulin (unlabelled 80%, Hilyte-647-tubulin 20%) diluted in the assay buffer (BRB80, 150 μM GMPCPP, 0.2% Tween20, 0.5 mg/ml Casein, 20 mM D-glucose, 0.22 mg/ml glucose oxidase and 20 ug/ml catalase) to 10 μl subtilisin-treated biotin-labelled microtubules (MT_Sub) immobilized on the coverslip by anti-biotin antibody. The reaction was allowed for 15 min at room temperature.

Alternatively, 8 μM of HiLyte647-tubulin were incubated with 200 nM PACRG-mNeonGreen either 200 nM FAP20-mRuby3, or with 200 nM PACRG-mNeonGreen and 200 nM FAP20-mRuby3 together for 15 min at RT. In each case, microtubule doublets formation was monitored under these conditions. Time-lapse image sequences were recorded for 15 min at the rate of 1 frame per 5 s with an exposure time 100 ms. Data from four independent experiments were collected, each experiment was repeated at least on three days.

### FRAP (fluorescent recovery after photobleaching)

Microtubule doublets (MTDs) were preassembled in the flow chamber by following *Microtubule doublet assembly assay* described previously, followed by flushing in mix of 250 nM PACRG-mNeonGreen together with 250 nM FAP20-mRuby3. The formation of HDRs and LDRs of both proteins were monitored. By using ROI sampler editor lines were drawn in the middle of regions of interest, defining future spot for belching. The 488-nM laser was used for photobleaching at 10% laser power for 1 s illumination with following imaging at triggered acquisition 488 nm - 561nm sequence: 30 s prebleach imaging at the rate of 1 frame per 5 s, bleach, 2 min recovery imaging at the rate of 1 frame per 2 s.

### Microtubule doublets dynamic assay

Microtubule doublets (MTDs) were preassembled by following *Microtubule doublet assembly assay* and HDRs were formed by flushing in 500 nM PACRG-mNeonGreen together with 500 nM FAP20-mRuby. To enhance a dynamic microtubule assay we subsequently flushed in the assay buffer (BRB80, 1 mM GTP, 0.2% Tween20, 0.5 mg/ml Casein, 20 mM D-glucose, 0.22 mg/ml glucose oxidase and 20 ug/ml catalase) with free 16μM HyLite-647 tubulin in the absence or presence of 500 nM FAP20-mRuby together with 500 nM PACRG-mNeonGreen (the mixture was pre-incubated on ice for 10 min to enhance proteins affinity to microtubules). Time-lapse image sequences were recorded for 20 min at the rate of 1 frame per 10 s with an exposure time 100 ms. Data from at least three independent experiments was collected.

### Microtubule doublets depolymerization assay

Microtubule doublets (MTDs) were preassembled by following *Microtubule doublet assembly assay* followed by growth of GDP-B-microtubule elongations in the presence of Taxol. For the B-MT elongations we used 6 uM free HyLite-647 tubulin supplemented with an assay buffer (BRB80, 1 mM GTP, 10μM taxol, 0.2% Tween20, 0.5 mg/ml Casein, 20 mM D-glucose, 0.22 mg/ml glucose oxidase and 20 ug/ml catalase). After 2 min 30 s of polymerisation, the process was stopped and elongation was stabilized by flushing out free tubulin by washing buffer with Taxol (BRB80, 10μM Taxol, 0.2% Tween20, 0.5 mg/ml Casein, 20 mM D-glucose, 0.22 mg/ml glucose oxidase and 20 ug/ml catalase). Further HDR were formed on GMPCPP B-MTs as well as on of GDP-B-microtubule elongations by flushing in 100 nM PACRG-mNeonGreen together with 100 nM FAP20-mRuby, both dissolved in the washing buffer with Taxol.

To study the depolymerization velocity of B-MTs we exchanged the washing buffer Taxol-free buffer (BRB80, 0.2% Tween20, 0.5 mg/ml Casein, 20 mM D-glucose, 0.22 mg/ml glucose oxidase and 20 ug/ml catalase) while maintaining the concentrations of both proteins in the solution. For the control experiment, all steps were repeated without the addition of any proteins. Time-lapse image sequences were recorded for 10 min at the rate of 1 frame per 5 s for *Microtubule doublet assembly assay,* for *Microtubule doublets depolymerization assay* time-lapse image sequences were recorded for 15 min at the rate of 1 frame per 5 s with an exposure time 100 ms. Data from three independent experiments was collected.

### In vitro binding of FAP20-mNeonGreen and PACRG-mNeonGreen to microtubule lattice with different post-translation modification

Mouse Taxol-wild-type microtubules were immobilized on anti-tubulin antibody treated channels. Then the channels were flushed with 40 μl of BRB80 to remove unbound microtubules and snapshot was taken to mark the position of wild type microtubules. Subsequently Taxol-HeLa-MTs or PTM knockdown Taxol-stabilized microtubules (*Ttll1^-/-^, Ttll7^-/-^*, *Ttll1^-/-^Ttll7^-/-^*, *Atat1^-/-^*) were immobilized and the channels were flushed with 40 μl of BRB80 to remove unbound microtubules and again the snapshot was taken to mark the position of microtubules. The same setup was used for tyrosinated and detyrosinated microtubules. Finally, the flow chamber was flushed with 20 μl of 50 nM PACRG-mNeonGreen or 50 nM FAP20-mNeonGreen diluted in the assay buffer (BRB80, 0.2% Tween20, 0.5 mg/ml Casein, 20 mM D-glucose, 0.22 mg/ml glucose oxidase and 20 ug/ml catalase). Time-lapse image sequences of PACRG-mNeonGreen or FAP20-mNeonGreen binding to microtubules were recorded for 1 min at the rate of 1 frame per 2 s with an exposure time 100 ms. Data from three independent experiments for each condition was collected.

## Image analysis

Microscopy data were analyzed using ImageJ (FIJI)^43^.

### HDRs and LDRs density estimation

High-Density Regions and Low-Density Regions of PACRG and FAP20 intensities on the assembled B-microtubules were measured in FIJI by drawing a rectangle around the B-MTs with HDR and B-MTs with LDR and measuring the average intensity per pixel separately for PACRG in green channel and for FAP20 in mCherry channel. The rectangle was then moved to an area directly adjacent to the B-microtubule where no microtubule is present and the average intensity per pixel was measured again and subtracted from of the average intensity per pixel the microtubule. The final densities of PACRG and FAP20 in the HDR were normalized to the average densities of PACRG and FAP20 in the LDR of B-MTs.

### Coverage of HDR on B-microtubules

Percentage of coverage of High-Density regions of PACRG and FAP20 was analyzed by measuring total length of B-MTs as well as total length of HDRs separately for PACRG and FAP20 per field of view using Kymographs. Kymographs were generated using KymographBuilder plug-in In FIJI by manually fitting lines along the A-and B-microtubules.

### FRAP (fluorescent recovery after photobleaching)

The intensity of bleached areas over time were measured In FIJI by manually fitting lines along the HDRs and LDRs of bleached areas and the intensities per pixel for PACRG-mNeonGreen and FAP20_mRuby separately were measured at every taken frame. The line was then moved to an area adjusted to the bleached regions where no microtubule is present and the intensity per pixel and per time frame was measured again and subtracted from the intensity per pixel of the bleached region for corresponding protein. The final densities of bleached region were normalized to the average densities of PACRG and FAP20 at the region before bleaching. Normalized data were fitted by using exponential recovery model:

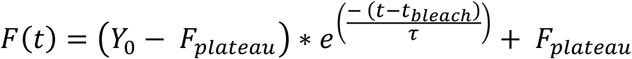

Where F(t) is the fluorescence intensity at time t. *Y*_0_ is the fluorescence intensity immediately after the bleach at *t_blech_. F_plateau_* is the fluorescence intensity after complete recovery, when fluorescence has stabilized and τ is recovery time constant. The immobile fraction (IF) for each protein in different condition was calculated using the formula

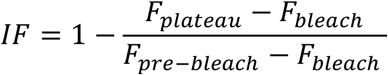

Where is *F_bleach_* is intensity immediately after blech and *F_pre−bleach_* the fluorescence intensity before bleach.

### Dynamic parameters of DMTs

Dynamic instability parameters of A-and B-microtubules as polymerization rate, depolymerization rate, frequency of catastrophes and frequency of rescues was analysed using Kymographs, which were generated using KymographBuilder plug-in In FIJI by manually fitting lines along the A-and B-microtubules.

### PACRG-mNeonGreen and FAP20-mNeonGreen density estimation

PACRG/FAP20 density on the microtubules with different post-translation modification was measured In FIJI by drawing a rectangle around the microtubule and measuring the average intensity per pixel. The rectangle was then moved to an area directly adjacent to the microtubule where no microtubule is present and the average intensity per pixel was measured again and subtracted from of the average intensity per pixel the microtubule. The final density of PACRG/FAP20 was normalized to the average of density PACRG/FAP20 on mouse wild type microtubules.

The same method was used for density estimation of PACRG-mNeonGreen and FAP20-mNeonGreen on detyrosinated and tyrosinated microtubules. The final density of PACRG/FAP20 was normalized to the average of density PACRG/FAP20 on tyrosinated microtubules.

### Cryo-EM grid preparation and cryo-tomograms acquisition

For control sample, MTDs in Nucleation Buffer (NB) were applied onto lacey carbon film grids (300 Mesh, Copper, EMS) for 30 seconds at 37°C and 90% humidity followed by 1s back blotting. 10 nm colloidal gold beads (EMS EM BSA Gold Tracer, cat number 25487) were applied to the grid, blotted for 1s and immediately vitrified in liquid ethane using Leica EM GP2 plunge freezer. For MTDs with FAP20 and PACRG, first MTDs in NB were applied onto the lacey carbon film grids (300 Mesh, Copper, EMS) for 30 seconds at 37°C and 90% humidity followed by 1s back blotting. FAP20 and PACRG were mixed in Flushing Buffer (FB) at 1 μM final concentration on ice to form the complex and applied on MTDs for 30s at 37°C and 90% humidity and blotted for 1s. Finally, 10 nm colloidal gold beads were applied and blotted for 1s and immediately vitrified in liquid ethane using Leica EM GP2 plunge freezer.

Images in cryo-electron microscopy were acquired with a Talos Arctica equipped with Falcon 4i detector camera and operating at 200 kV. Tilt-series were collected automatically from-60° to +60° with 3° angular increment at 63000X of magnification (pixel size = 1.96 Å), using a defocus range of-3 μm to-5 μm for a total dose of 40 e-/Å2. The cryo-electron tomograms were aligned and reconstructed using IMOD using gold beads as fiducial markers. A total of 62 tomograms of control MTDs and 53 tomograms of FAP20 and PACR3 incubated MTDs were acquired. Reconstructed tomograms were binned four times (pixel size = 7.892) and denoised using IsoNet for representative images in Fig. 5 (Fig 5 A, B, C and F, G, H). Images presented in Figure 5A and 5F, correspond to projection averages of 50 and 80 tomogram slices, respectively, using ZProjection tool on Fiji software^43^.

### Curvature of B-microtubule

To study the curvature of the B-microtubules, a set of 46 B-microtubules have been selected from 23 tomograms for control MTDs and 40 B-microtubules have been selected from 22 tomograms for MTDs with FAP20 and PACRG. The tomograms were resliced from the top and projected with average intensity using grouped z-projection method with a group size of 10 image stacks. The curvature was measured at different positions along these doublets. Each time, several coordinates were picked using the point picker plug-in on Fiji software to describe the shape of the B-microtubule: the center of the A-microtubule (O), the position on the A-microtubule where B-microtubule was attached (OJ) and six points on each B-microtubule. In total, 1344 and 1226 MTD sections were measured for control and FAP20-PACRG, respectively. The same picking was done for the MTD in vivo using the map EMD 8528.

To superimpose these sets together, the segment O-OJ was used as a reference. Each set of coordinates was rotated and shifted to have all reference points aligned using a homemade R script (https://www.R-project.org/ and https://github.com/CentrioleLab/MTD_curvature). In Fig. 5, a representative plot was shown for the corresponding doublet as well as the total of picked coordinates for both conditions with the overall averages (Figure5 D, E, I, J). To highlight the flexibility or the stability of the B-microtubule, three schematic doublets were generated; one with in vivo curvature only, and the others with the in vitro curvatures for two conditions (blue or orange lines). All three images were open in Fiji to generate a plot profile following the dashed arrow lines (1 and 2). The plot profiles of control MTDs and MTDs with FAP20 and PACRG were superimposed with the peak of in vivo curvature (vertical black line) (Related to Fig5 K-N).

---

**Fig S1.**
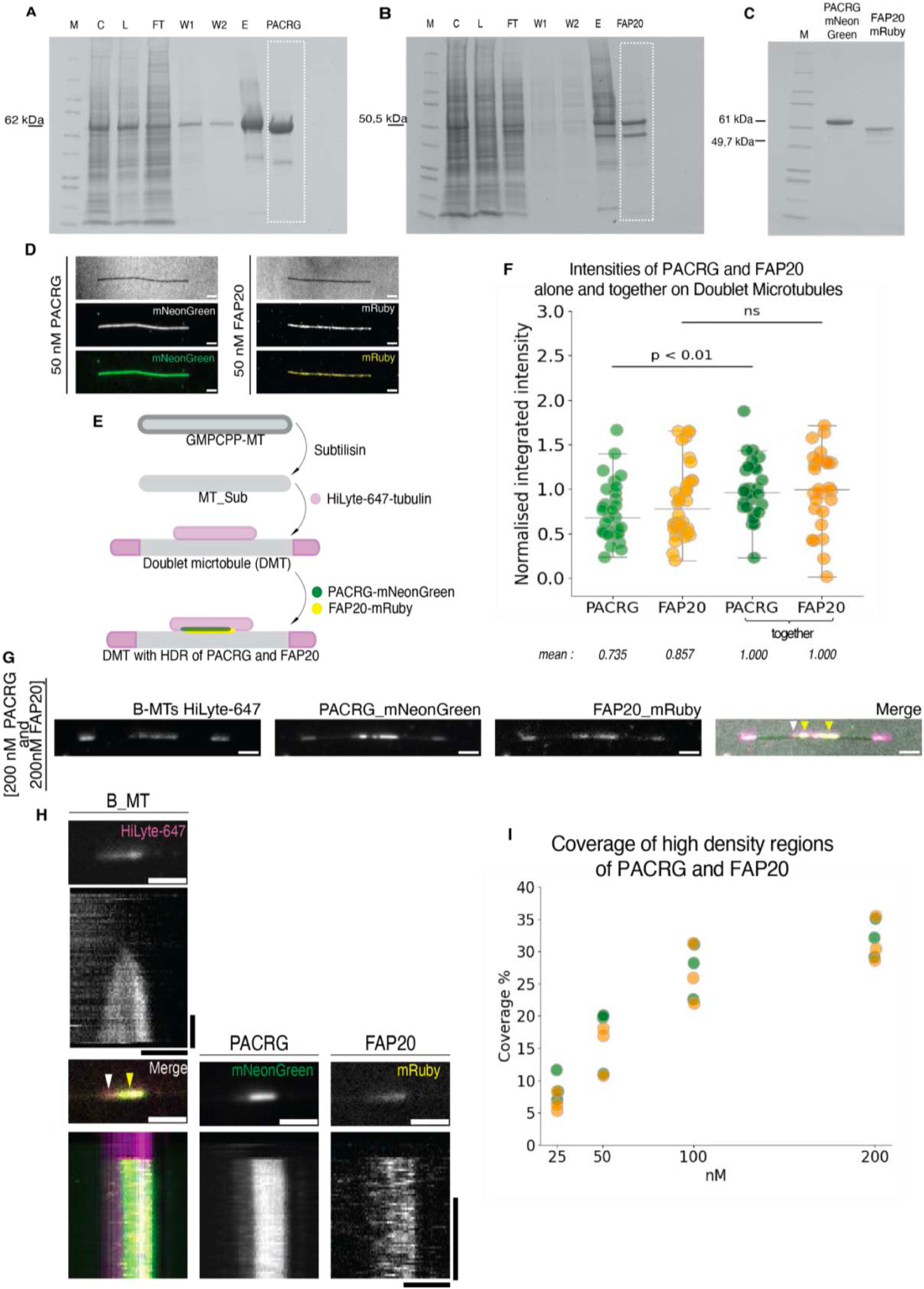
(A) SDS-PAGE gel of Ni-NTA affinity purification of PACRG-mNeonGreen. M-Precision Plus Protein Standard, Kaleidoscope. C – cells resuspended in lysis buffer. L – lysate. FT – flow through. W1, W2 – first and second washing fraction. E – elution of PACRG-mNeonGreen-His, Mw = 62 kDa. PACRG – PACRG-mNeonGreen with cleaved His-tag from N-terminus, Mw = 61 kDa. White dashed square indicates PACRG-mNeonGreen fraction before SEC. **(B)** SDS-PAGE gel of Ni-NTA affinity purification of FAP20-mRuby. M-Precision Plus Protein Standard, Kaleidoscope. C – cells resuspended in lysis buffer. L – lysate. FT – flow through. W1, W2 – first and second washing fraction. E – elution of FAP20-mRuby-His, Mw = 50,5 kDa. FAP20-mRuby with cleaved His-tag from N-terminus, Mw = 49,7 kDa. White dashed square indicates FAP20-mRuby fraction before SEC. **(C)** SDS-PAGE gel of SEC of FAP20-mRuby and PACRG-mNeonGreen. M-Precision Plus Protein Standard, Kaleidoscope. FAP20 - FAP20-mRuby Mw =49,7 kDa. PACRG - PACRG-mNeonGreen Mw =61 kDa. **(D)** Binding of 50 nM PACRG-mNeonGreen, depicted in green and FAP20-mRuby, depicted in yellow to GMPCPP stabilized microtubules with C-terminal tails. Scale bar 2 μm. **(E)** Schematic drawing of microtubule doublets (MTDs) assembly and the formation of high-density region (HDR) by adding FAP20 and PACRG together to B-tubule (B-MT). **(F)** Normalized integrated intensities of PACRG and FAP20 in the co-assembly assay alone and in the low-density regions (LDRs), in the conditions when both proteins were present. p < 0.01, determined by the two-tailed t-tests Mann-Whitney U Test between PACRG alone and PACRG in the low-density regions LDRs. **(G)** Montage showing microtubule-doublets (MTDs) with high-density region (HDR) of PACRG and FAP20. A-tubule tips (white arrowheads) and B-tubules formed by 647-HiLyte tubulin are depicted in magenta. Yellow arrowheads indicate HDRs formed in the presence of 200 nM PACRG labeled by mNeonGreen together with 200 nM FAP20 labeled by mRuby. Scale bar, 2 μm. **(H)** Montage showing B-microtubule (B-MT) assembly without additional proteins and the formation of high-density region (HDR) of FAP20 and PACRG and the corresponding kymograph. Horizontal scale bar, 2 μm, vertical scale bar is 2 min. **(I)** Percentage of high-density region (HDR) coverage of FAP20 (orange) and PACRG (green) on B-tubules (B-MTs). Groups represent HDRs formation in the presence of PACRG and FAP20 together on different concentration range 25 nM, 50 nM, 100 nM and 200 nM (N=3, number of independent experiments for each condition).

**Fig S3.**
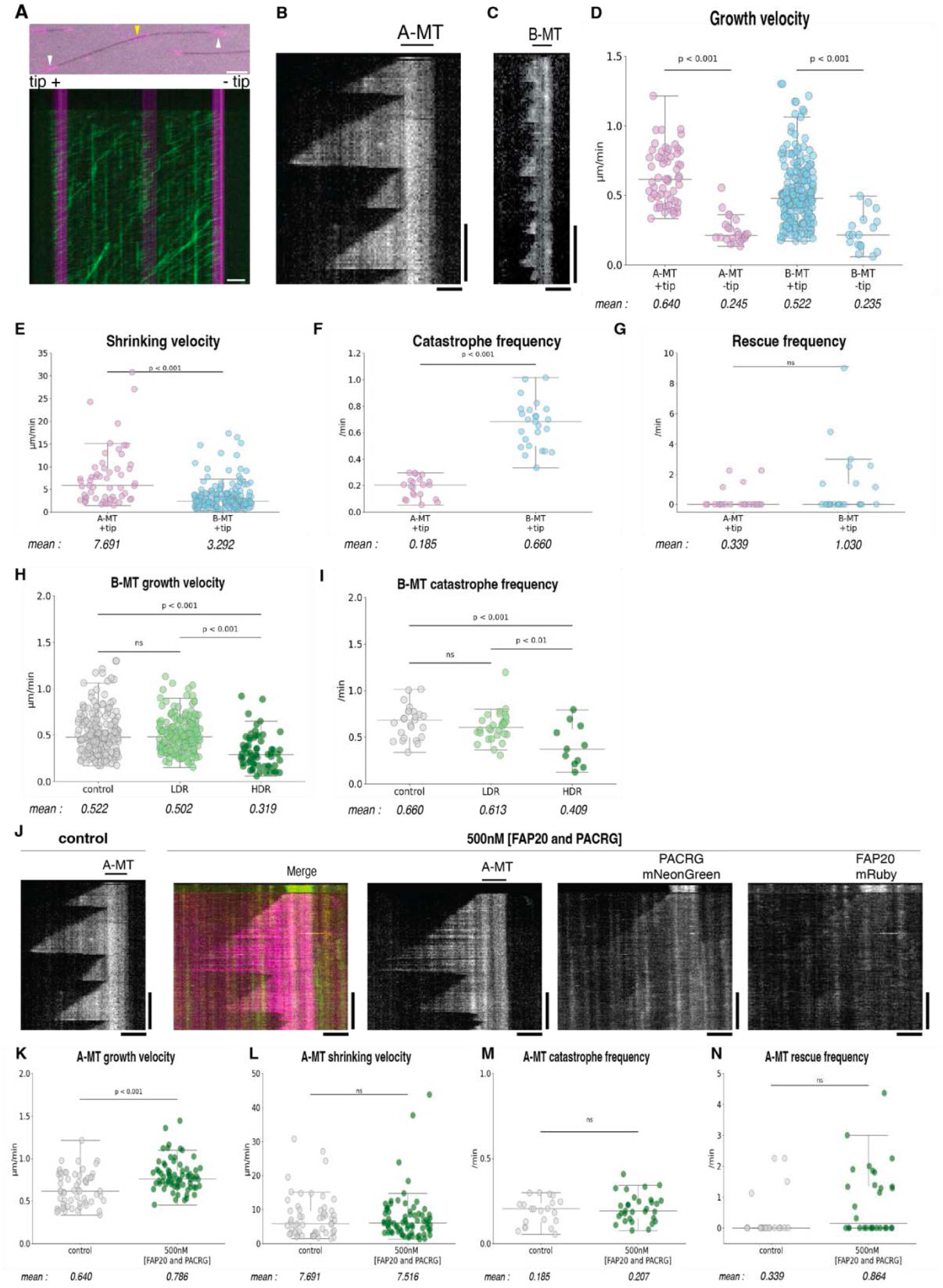
The dynamic of A-and B-tubules. (**A**) Montage showing pre-assembled B-tubule and corresponding kymograph of walking short Kif5B in one direction towards plus end. (**B**) Montage showing kymographs of dynamic A-and (**C**) B-tubules. Scale bar, horizontal-2 μm and vertical - 5 min. (**D**) Growth velocity of A-tubule (A-MT, depicted in pink) and B-tubule (B-MT, depicted in blue) at +tip and-tip. (**E**) Shrinking velocity of A-tubule (depicted in pink) and B-tubule (depicted in blue) at +tip. (**F**) Catastrophe frequency of A-tubule (depicted in pink) and B-tubule (depicted in blue) at +tip. (**G**) Rescue frequency of A-tubule (depicted in pink) and B-tubule (depicted in blue) at +tip. (**D**-**G**) P-values, determined by the two-tailed t-test. (**H**) Growth velocity (**I**) Catastrophe frequency (**H**-**I**) of B-tubules without additional proteins (control-depicted in grey), with low-density binding of proteins (LDR-depicted in light green), and with high-density binding of proteins (HDRs - depicted in dark green). Final concentration of proteins in the solution 500nM PACRG-mNeonGreen and 500 nM FAP20-mRuby3. P-values, determined by the two-tailed t-test. (**J**) Montage showing kymographs of dynamic A-tubules without and with 500mM PACRG-mNeonGreen and 500 nM FAP20-mRuby. Scale bar, horizontal - 2 μm and vertical - 5 min. (**K**-**N**) Dynamic parameters of A-microtubules without and with 500mM PACRG-mNeonGreen and 500 nM FAP20-mRuby: (K) growth velocity; (L) shrinking velocity; (M) catastrophe frequency; (N) rescue frequency. Control-depicted in grey, dynamic of A-tubules in the presence of proteins - depicted in dark green. P-values, determined by the two-tailed t-test.

**Fig S4.**
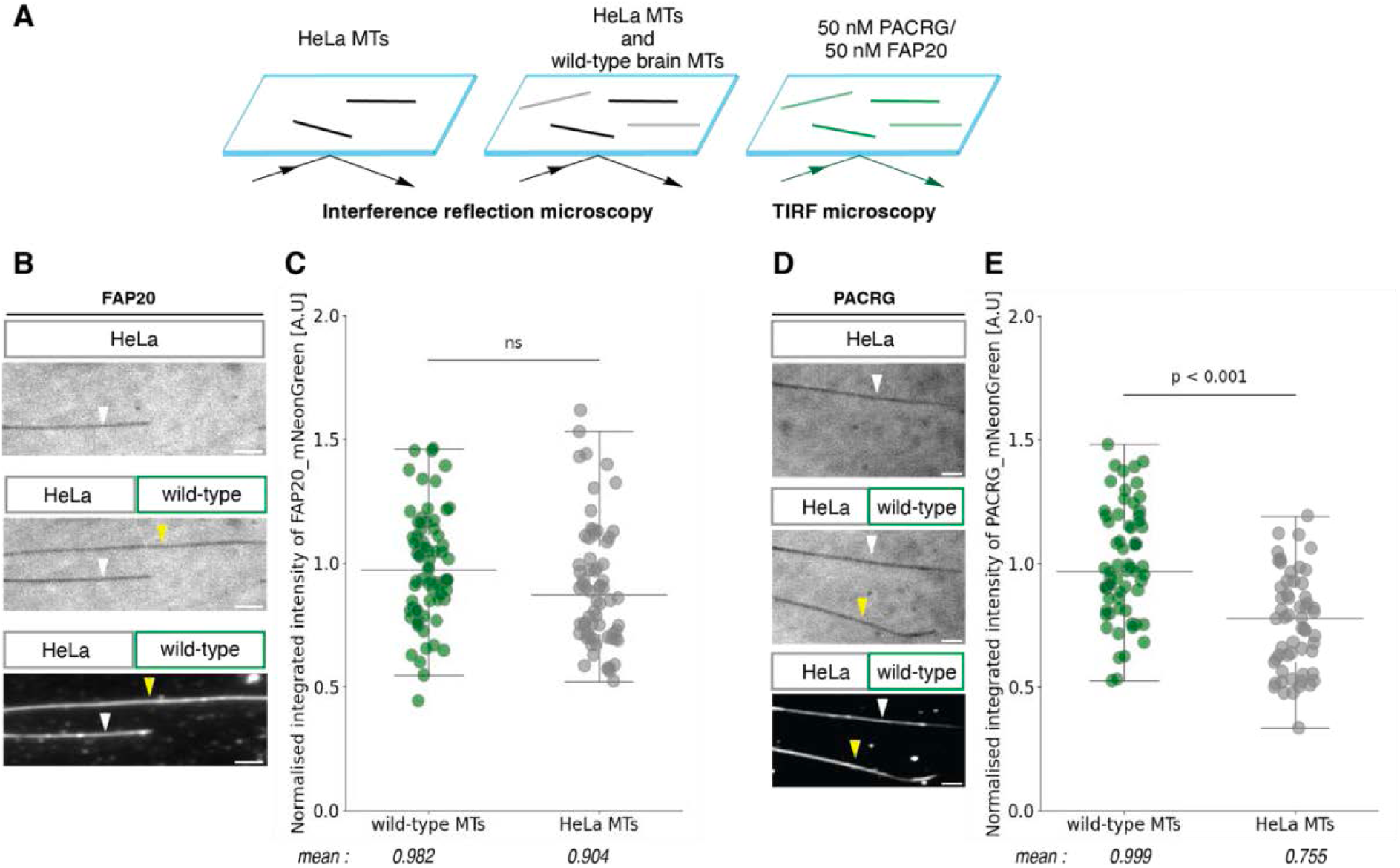
Binding of PACRG-mNeonGreen and FAP20-mNeonGreen to HeLa and wild-type brain microtubules. (**A**) Schematic drawing of the assay setup. (**B, D**) Montage showing IRM and TIRF microscopy images of the HeLa and wild-type brain microtubules and intensities of FAP20-mNeonGreen and PACRG-mNeonGreen respectively. White arrowheads indicate HeLa microtubules, yellow arrowheads – wild type brain microtubules. Scale bar, 2μm. **(C, E)** Normalized integrated average intensities of PACRG-mNeonGreen and FAP20-mNeonGreenon respectively on taxol-microtubules, in three independent sets of the experiment N=3.P-values, determined by the Mann-Whitney U Test.

